# SARS-CoV-2 epitope mapping on microarrays highlights strong immune-response to N protein region

**DOI:** 10.1101/2020.11.09.374082

**Authors:** Angelo Musicò, Roberto Frigerio, Alessandro Mussida, Luisa Barzon, Alessandro Sinigaglia, Silvia Riccetti, Federico Gobbi, Chiara Piubelli, Greta Bergamaschi, Marcella Chiari, Alessandro Gori, Marina Cretich

## Abstract

A workflow for SARS-CoV-2 epitope discovery on peptide microarrays is herein reported. The process started with a proteome-wide screening of immunoreactivity based on the use of a high-density microarray followed by a refinement and validation phase on a restricted panel of probes using microarrays with tailored peptide immobilization through a click-based strategy. Progressively larger, independent cohorts of Covid-19 positive sera were tested in the refinement processes, leading to the identification of immunodominant regions on SARS-CoV-2 Spike (S), Nucleocapsid (N) protein and Orf1ab polyprotein. A summary study testing 50 serum samples highlighted an epitope of the N protein (region 155-171) providing 92% sensitivity and 100% specificity of IgG detection in Covid-19 samples thus being a promising candidate for rapid implementation in serological tests.

## Introduction

In the frame of the ongoing pandemic caused by Severe Acute Respiratory Syndrome Coronavirus 2 (SARS-CoV-2), reliable diagnostic and prognostic tools are constantly needed for the appropriate management of patients, to guide the development and implementation of surveillance and prevention measures, and to assess the efficacy of therapeutic interventions and vaccines. While molecular and antigenic testing of respiratory samples are used for the diagnosis of acute SARS-CoV-2 infection, serology testing allows the identification of individuals with recent or past exposure to SARS-CoV-2. Serological tests can be combined with molecular testing to improve the sensitivity and specificity of diagnosis^1^ and represent key instruments for epidemiological surveillance to guide Public Health interventions ^2^. Data from representative and reliable seroprevalence studies will greatly support decision-making practices in terms of physical distancing and other restrictions. However, the use of seroprevalence data to inform policymaking strictly depends on the accuracy and reliability of tests. Yet, currently available serological assays are biased by false-positive and false-negative results as pointed out by sub-optimal sensitivity and specificity performance (https://www.fda.gov/medical-devices/emergency-situations-medical-devices/eua-authorized-serology-test-performance).

To date, the majority of the serological studies confirmed a general humoral response upon SARS-CoV-2 infection against spike (S) and nucleocapsid (N) proteins, observed as early as the 4th day after symptom onset, when SARS-CoV-2-specific IgG and IgM antibodies appear simultaneously or sequentially and plateau within 6 days after seroconversion ^3, 4^. Most current immunoassays use recombinant coronavirus proteins: spike (S), which is exposed on the surface of the viral particle, and nucleocapsid (N), which is highly expressed during infection ^5, 3^.

Full-length recombinant antigens, however, in addition to high costs and storage constraints, may suffer from stability problems, batch-to-batch variations and in some cases may give rise to cross-reactivity leading to ambiguous detection outcomes. This might be the case for SARS-CoV-2 when using proteins sharing high homology with genetically similar, co-circulating, human coronaviruses (hCoV) that are responsible for common cold ^6, 7^.

As a further element, a fine dissection of the specific regions on the viral proteins recognized by virus-specific antibodies have yet to be fully characterized. This knowledge could improve our understanding of diverse disease outcomes and inform the development of improved diagnostics, vaccines, and antibody-based therapies ^8^

In this scenario, peptide microarrays are appealing tools as they allow the simultaneous screening of hundreds to thousands different peptides immobilized in spatially ordered patterns of spots on solid supports ^9 10^ possibly leading to the development of synthetic diagnostic probes with improved specificity and of putative, new immunotherapeutic targets.

Herein we report on SARS-CoV-2 linear epitopes identified by probing a proteome-wide peptide microarray with serum samples collected from seven elderly patients with mild/moderate COVID-19 about one month after symptom onset. The epitope mapping resulted in the selection of 12 putative linear epitopes located on the SARS-CoV-2 N protein, S protein, and ORF1ab polyprotein.

Taking advantage of our previously developed microarray platform for site-selective and oriented peptide immobilization ^11^, we validated the 12 selected probes with sera collected from an independent group of twelve COVID-19 patients at about one month after symptom onset and in a group of pre-COVID-19 serum samples collected before December 2018 ^11^(Scheme 1). The best performing linear epitope turned out to belong to the N protein (region 155-171), which was further validated with serum samples collected from 50 COVID-19 patients at 1-5 months after symptom onset, including 5 patients with sera collected at different time-points after symptom onset.

**Scheme 1:**
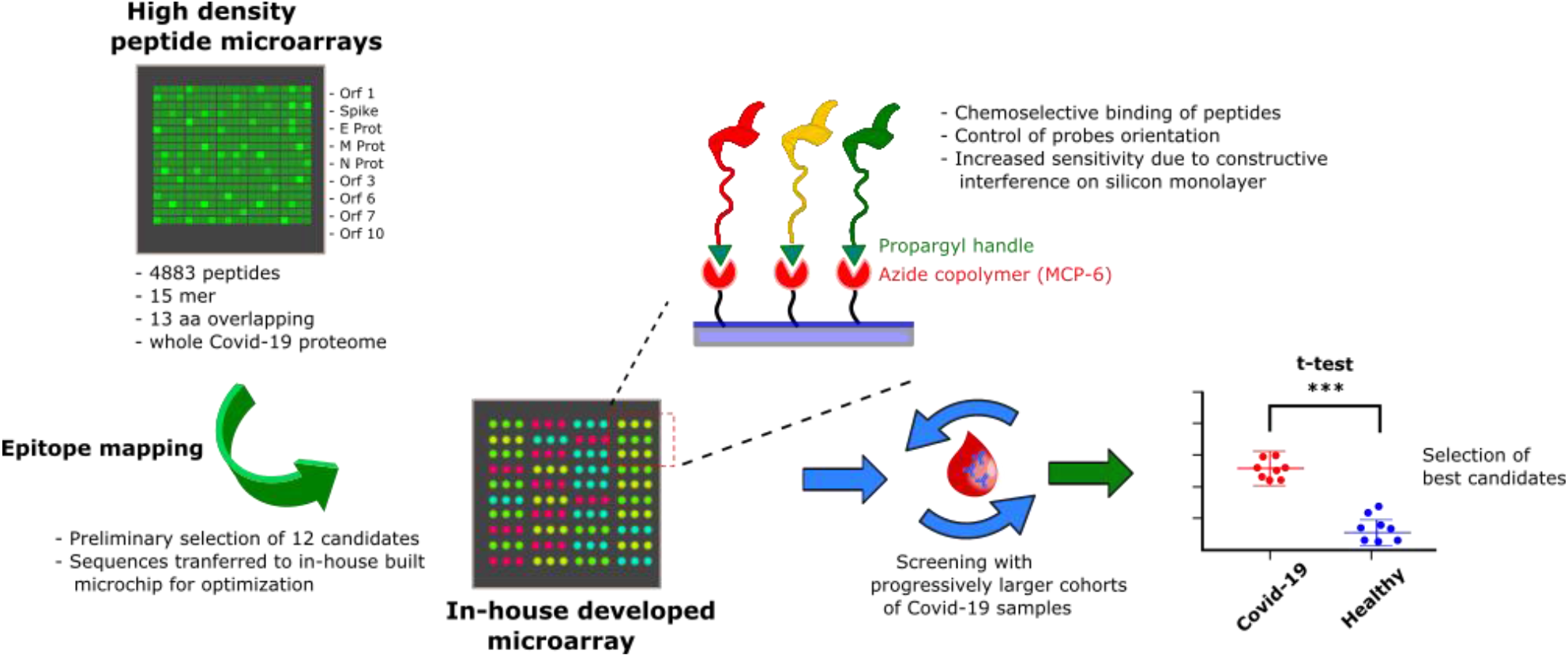
Overview of the probe selection workflow. A high-density peptide microarray displaying the whole SARS-CoV-2 proteome was probed for immunoreactivity with COVID-19 serum samples leading to the selection of 12 peptide hits. The most promising candidates were then transferred to a low-density microarray platform for site-selective and oriented peptide immobilization and validated with progressively larger and independent cohorts of patients to select the most sensitive and specific immunoreactive peptide probes.

## Results and discussion

### SARS-CoV-2 whole-proteome epitope mapping

Preliminary data on IgG response in sera of COVID-19 patients were generated using high-density peptide array made of 4883 different peptides, 15 amino acids long, 13 amino acids overlap (PEPperCHIP® SARS-CoV-2 Proteome Microarray) displaying sequences of ORF1ab polyprotein, surface glycoprotein (S), ORF3a protein, envelope protein (E), membrane glycoprotein (M), ORF6 protein, ORF7a protein, ORF8 protein, nucleocapsid phosphoprotein (N) and ORF10. Serum samples were collected from a group of seven COVID-19 patients, aged 68-96 years, at about 30 days after the onset of symptoms. In these patients, who had mild or moderate COVID-19, diagnosis of SARS-CoV-2 infection was confirmed by molecular testing of nasopharyngeal swabs and the presence of SARS-CoV-2-specific antibodies in corresponding serum samples was demonstrated by virus microneutralization assay. Table 1 reports demographic and clinical data of the COVID-19 patients tested in this study.

**Table 1:**
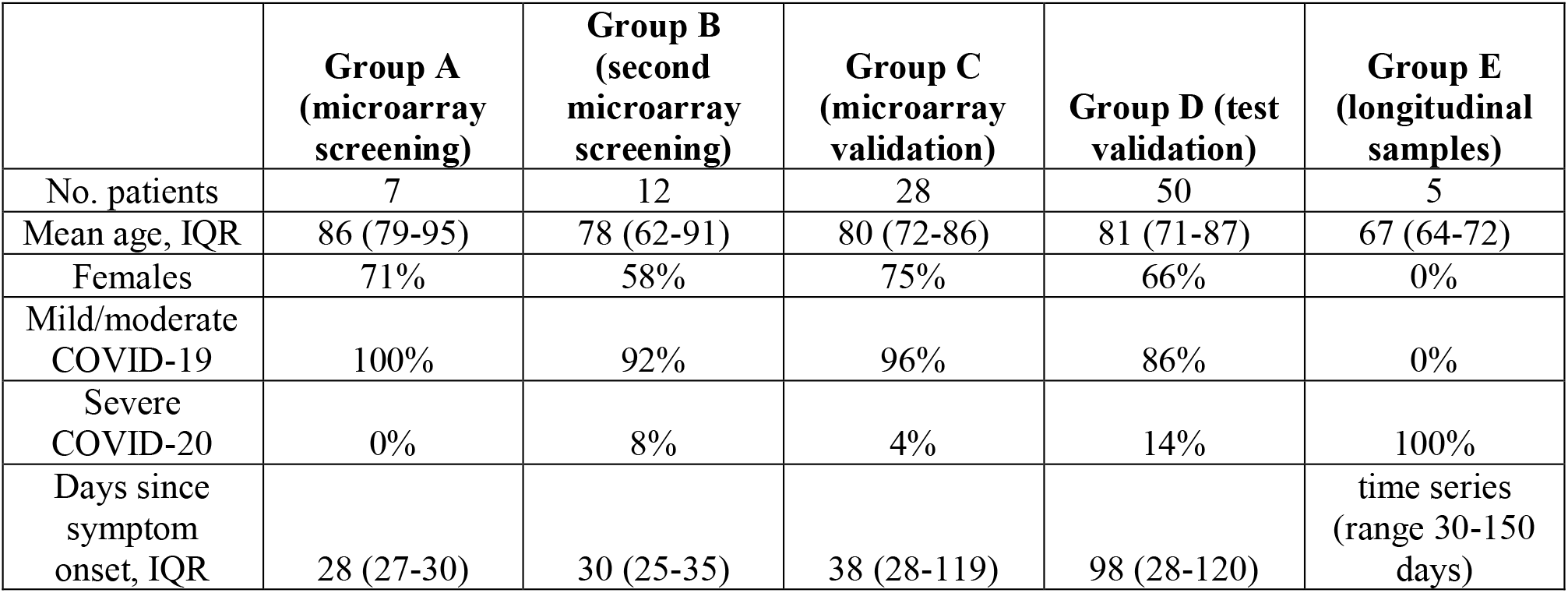
demographic and clinical data on the COVID-19 patients tested.

|  | **Group A**<br>**(microarray**<br>**screening)** | **Group B**<br>**(second**<br>**microarray**<br>**screening)** | **Group C**<br>**(microarray**<br>**validation)** | **Group D (test**<br>**validation)** | **Group E**<br>**(longitudinal**<br>**samples)** |
| --- | --- | --- | --- | --- | --- |
| No. patients | 7 | 12 | 28 | 50 | 5 |
| Mean age, IQR | 86 (79-95) | 78 (62-91) | 80 (72-86) | 81 (71-87) | 67 (64-72) |
| Females | 71% | 58% | 75% | 66% | 0% |
| Mild/moderate<br>COVID-19 | 100% | 92% | 96% | 86% | 0% |
| Severe<br>COVID-20 | 0% | 8% | 4% | 14% | 100% |
| Days since<br>symptom<br>onset, IQR | 28 (27-30) | 30 (25-35) | 38 (28-119) | 98 (28-120) | time series<br>(range 30-150<br>days) |

The whole-proteome peptide microarray screening was performed according to manufacturer instructions after serum samples inactivation at 56°C for 30 minutes (see Methods).

Epitope mapping results were analyzed in terms of fluorescence intensity of IgG immune-reactivity of COVID-19 positive sera and discrimination with the healthy control group by the unpaired *t* test (significative p<0.05). Several immunoreactive regions were identified on nucleocapsid (N) protein, spike (S) protein and ORF1ab polyprotein. No reactivity was observed on the sequences from the other proteins displayed on the microarray. To define a set of immunodominant epitopes for further characterization, we selected the 10 most reactive antigenic regions corresponding to at least three contiguous overlapping peptides that showed a fluorescence intensity statistically different from the one detected in healthy controls. Two more immunoreactive sequences fulfilling the discrimination criteria were added to the previously selected, even if fluorescence was detectable only on a single sequence and not in the adjacent overlapping peptides. The selection of the 12 most reactive aminoacidic sequences is reported in Table 2; the resulting linear epitopes for protein S and N are mapped within antigen structures in Figure 1A.

**Table 2:**
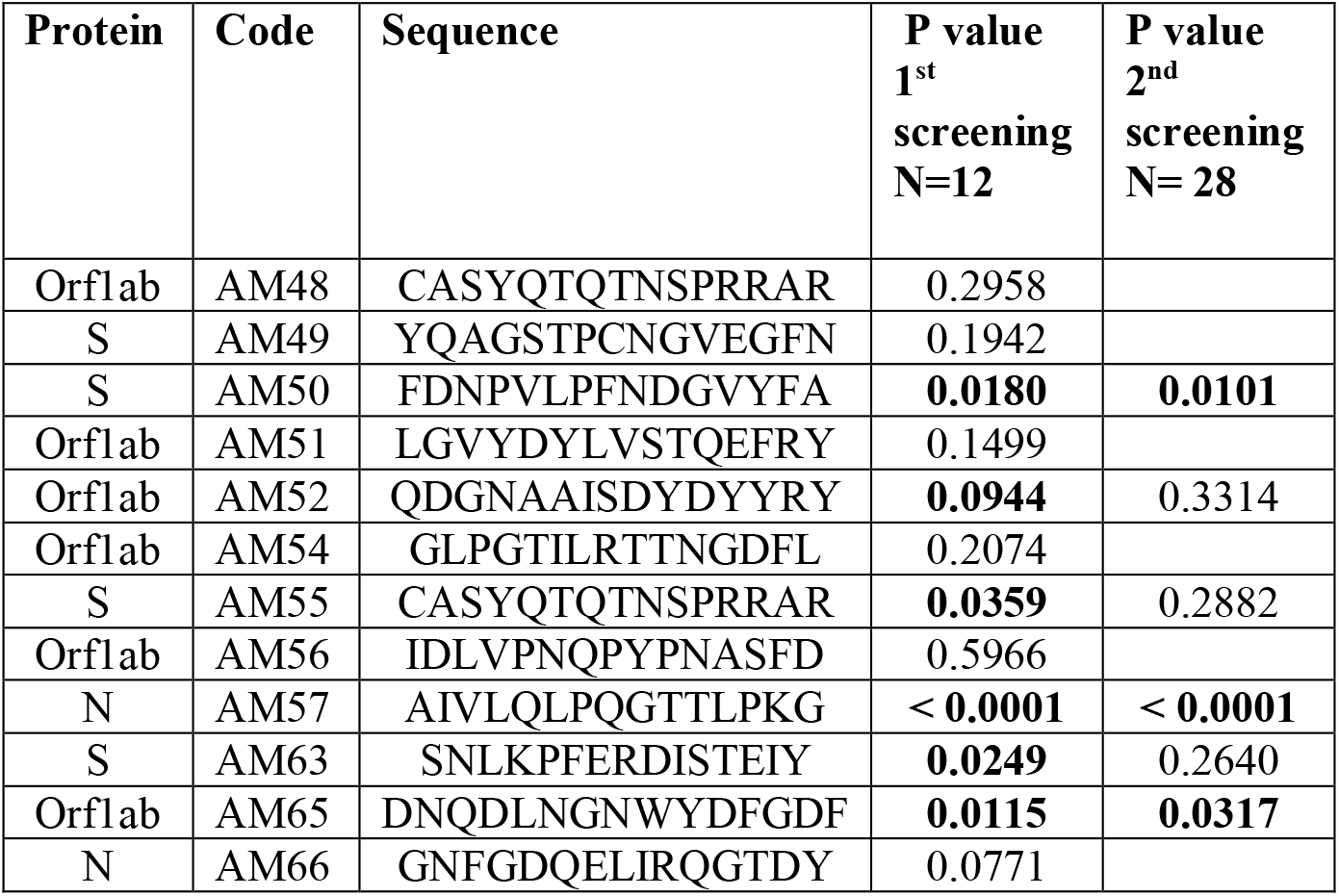
Most reactive aminoacidic sequences selected by the high-density microarray and results of the validation performed with progressively larger patients cohorts. The peptide sequences were analyzed in terms of fluorescence intensity of IgG immune-reactivity of COVID-19 positive sera and discrimination with the healthy control group by the unpaired *t* test. The p values of the screenings are reported

**Figure 1.**
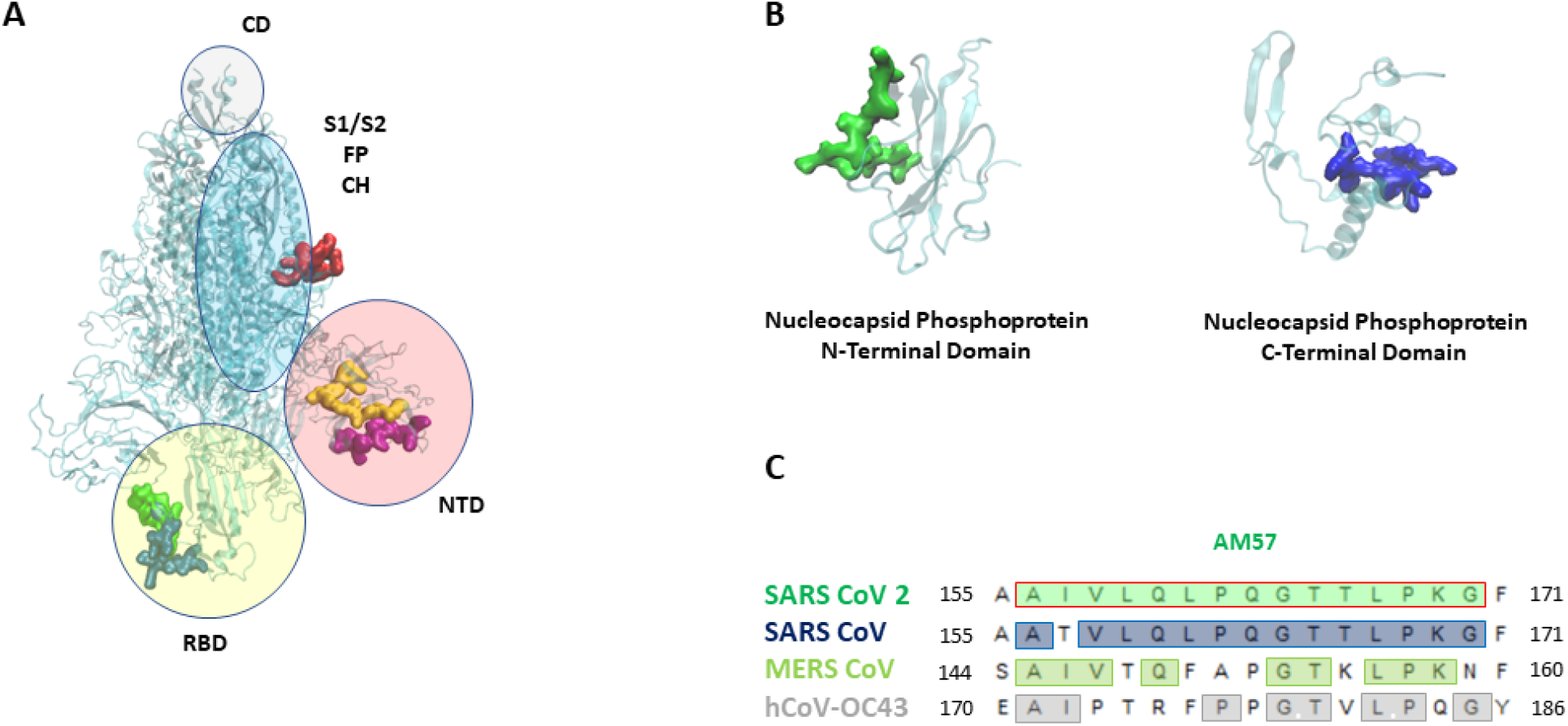
(A): SARS-CoV 2 Spike (S) protein with its domains ^12^ (NTD: n terminal domain, S1 e S2: furin cleavage site, CD: connecting domain, HR2: heptad repeat 2, FP: fusion peptide, CH: central helix). Epitopes are highlighted as follows: AM55 in red; AM50 in yellow; AM64 in purple; AM 63 in blue, AM 49 in green. (B): SARS-CoV 2 nucleocapsid (N) protein, the N-terminal and C-terminal domains. Epitopes are highlighted as follows: AM57 in green, AM66 in blue. (C): AM57 sequence alignment on SARS CoV 2, SARS CoV, MERS CoV, hCoV-OC43.

Notably, our data on SARS-CoV-2 proteins primarily targeted by COVID-19 patient antibodies (proteins S, N and ORF1ab) are in accordance with a detailed exploration of the humoral response in COVID-19 sera recently performed by a programmable phage-display immunoprecipitation and sequencing technology ^8^

### Epitopes validation screening

More focused validation screenings on the 12 hit epitopes derived from the high-density microarray were then performed by progressively increasing the number of patients tested and using independent cohorts. In these experiments, peptides were in-house synthesized including an azide handle, in order to apply a ‘click’ chemoselective strategy for peptide binding to polymer-modified microarrays slides (Scheme 1). This strategy was previously shown to allow for fine control of ligand orientation and probe surface density, which correlated with improved assay performances ^12–14^. The initial set of 12 probes was first screened for IgG immunoreactivity against sera collected from a cohort of N =12 COVID-19 patients at about 30 days after symptom onset *vs* N =12 pre-COVID-19 controls (Figure 2) and their discrimination performance evaluated by the unpaired t test (Table 2). Six out of the twelve putative epitopes identified by the initial screening showed ability to discriminate the two sample groups at a statistically relevant level: AM50, AM55 and AM63 from the S protein, AM52 and AM65 form Orf1ab polyprotein and AM57 form the N protein.

**Figure 2:**
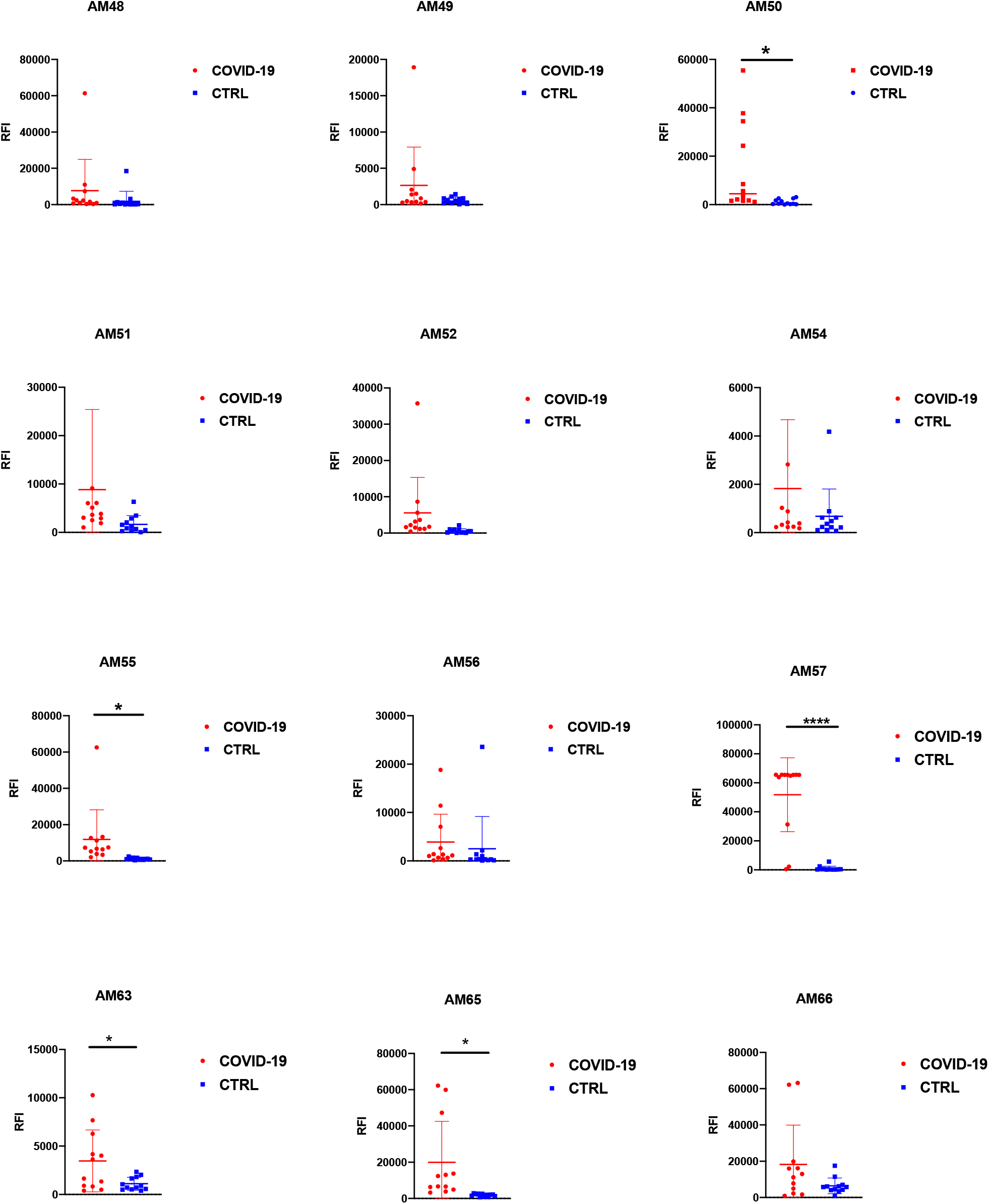
unpaired t-Test results for the peptide specific IgG. Peptide arrays displaying twelve peptide probes were probed with sera (N = 12) from COVID-19 patients and control patients (N = 12). Significative: p<0.05; * = p<0.05; ** = p<0.01; *** = p<0.001; **** = p<0.0001

The 6 sequences providing a p value <0.05 were re-evaluated in a second screening using a larger independent cohort of N = 28 patients, who were evaluated at 1-5 months after symptom onset (Figure 1 Supplementary Information). Again, testing a different and larger cohort of samples reduced the number of hit epitopes because a statistically significant ability to discriminate COVID-19 patients was found only in 3 out of 6 sequences: AM50 (S protein), AM65 (Orf1ab polyprotein) and AM57 located on the N protein whereas a loss of discrimination ability for probes AM52, AM55 and AM63 was observed (Table 2).

Epitope AM57 from the N protein consistently showed a high immunoreactivity, negligible response in healthy controls along all the screenings and the best diagnostic performance (p value <0.0001). We also performed alignment of AM57 sequence against SARS-CoV, MERS-CoV and hCoV-OC43 (www.bioinformatics.org), (Figure 1C), to assess whether it is conserved among the same virus family. Importantly, in light of possible pre-existing cross-reactive responses in healthy controls, AM57 (Figure 1-C) revealed only 53% amino acid identity with the corresponding sequence from the N protein of hCoV – OC43, responsible for the common cold.

The results of the study on N = 50 COVID-19 serum samples *vs* N=30 healthy controls of the IgG and IgM reactivity to the immunodominant AM57 epitope on N protein are summarized in Figure 3. Overall, for IgG (Figure 3, upper panel), AM57 showed an excellent diagnostic performance (p value <0.0001) in discriminating the two tested groups, with 92% sensitivity and 100% specificity, almost matching the performances given by the full N antigen (see Supplementary Information, Figure 2). After ROC curve analysis, AUC is 0.9747. The lack of false negative results detected among the N =50 COVID-19 samples suggests the absence of pre-existing cross-reactive response, likely due to low conservation of AM57 sequence among the common cold coronaviruses.

**Figure 3.**
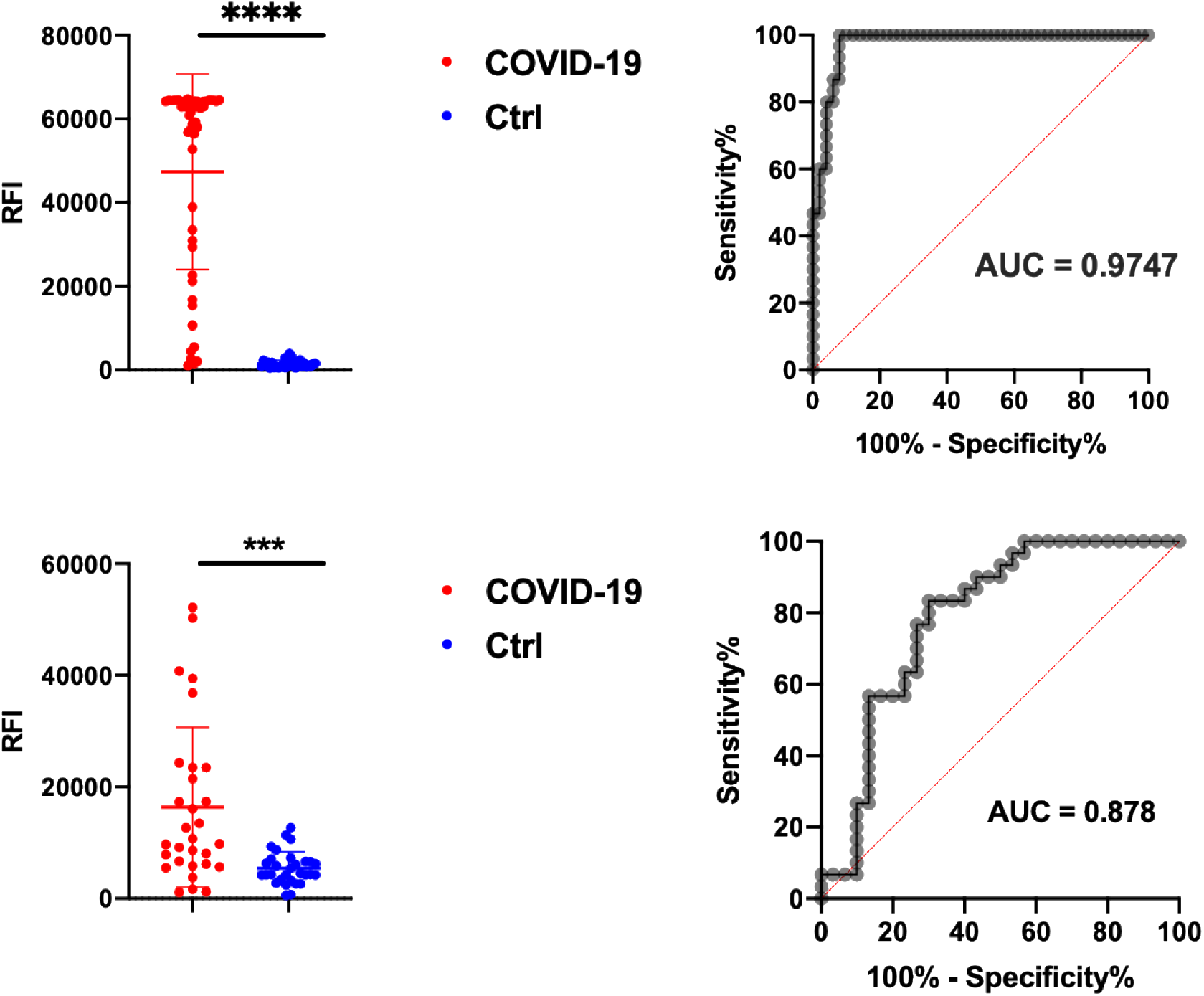
Upper panel: unpaired t-Test results for the peptide specific IgG to AM57 probed with sera (N = 50) from COVID-19 patients and control patients (N = 50) and corresponding AUC. Lower panel: unpaired t-Test results for the peptide specific IgM to AM57 probed with sera (N = 30) from COVID-19 patients and control patients (N = 30) and corresponding AUC. Significative: p<0.05; * = p<0.05; ** = p<0.01; *** = p<0.001; **** = p<0.0001

**Figure 4:**
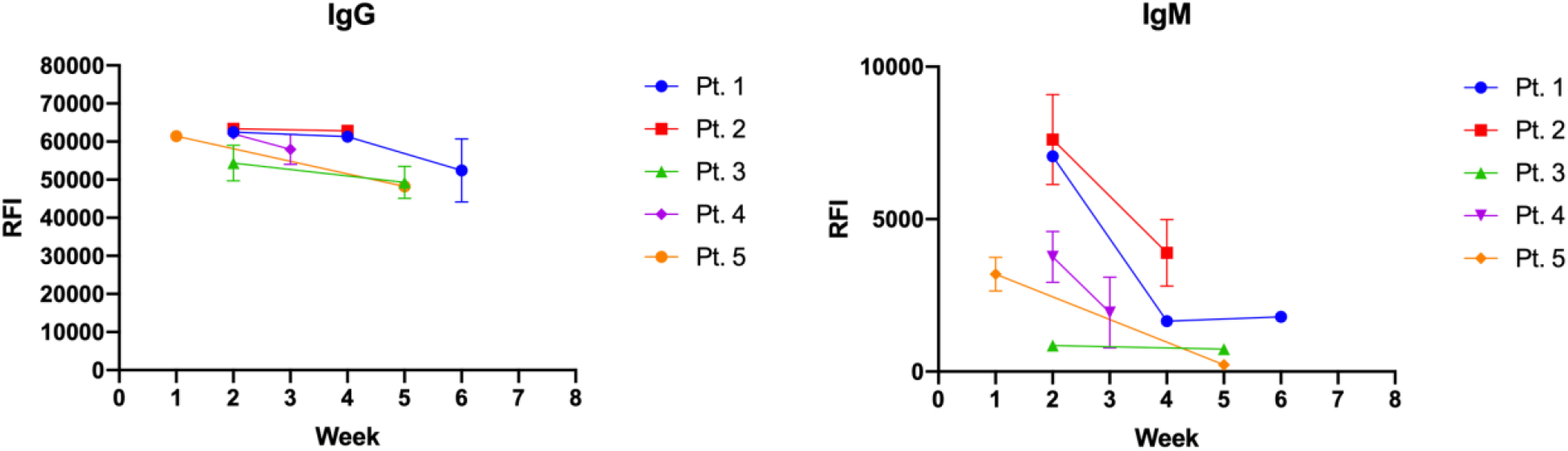
longitudinal immunoareactivity to AM57 of IgG (left panel) and IgM (right panel) in a set of 5 patients over a period of 6 weeks.

Also SARS-CoV-2 IgM immunoreactivity, which was tested in a set of N = 30 serum samples collected within the first month after symptom onset, showed a good discrimination ability (p value <0.001) and AUC = 0.878. (Figure 3, lower panel). Notably, the same test performed on the entire N antigen (See SI) confirmed a high variability of the IgM response among individuals.

To evaluate the longitudinal IgG and IgM profile on AM57, immunoreactivity at different timepoints following positivity to SARS-CoV-2 PCR test was evaluated (Figure 5) in 5 patients. While the IgG response showed to be quite stable over a period of 7 weeks, as expected the IgM response showed a decline. Both IgG and IgM immune-responses were detectable at two weeks since the molecular diagnosis.

Overall, the ability of the AM57 peptide to early detect IgM response worth further investigation, in line with the increasing interest on the use of peptide probes for point of care tests. The use of peptidic probes in place of full antigens is particularly appealing not only for storage stability, batch-to batch consistencies and avoidance of protein refolding issues, but also for the economic sustainability of population wide testing during a pandemics. Given the quickly spread around the globe, it is anticipated that rapid serological assays for COVID-19 will be needed by an increasing number of people and over a long period of time. In this frame, synthetic peptides offer clear advantages in terms of costs and synthetic versatility allowing straightforward implementation in diverse diagnostic settings: from lateral-flow-tests for point of need diagnostics to ELISA and bead-based assays for centralized settings.

## Conclusions

We here reported on a workflow for SARS-CoV-2 epitope discovery based on peptide microarrays. The discovery process started with a proteome-wide screening on high-density microarray followed by a refinement and validation phase based on low-density arrays characterized by a precise control of orientation and density of peptide probes. The results of the validation screenings showed the crucial role of testing a large number of samples and independent cohorts during a diagnostic probe refinement processes. This is particularly evident when looking at the high variability in immunoreactivity that is observable among the different sera in each set of sample. Our approach led to the identification of immunodominant regions on SARS-CoV-2 Spike, Nucleocapsid protein and Orf1ab polyprotein. A summary study testing 50 Covid-19 serum samples highlighted the optimal diagnostic performance of a N protein linear epitope (region 155-171) which can be considered a promising candidate for rapid implementation in cost-effective serological tests, thus expanding the tool box needed to complement Covid-19 patient assessment, vaccine research and epidemiological studies.

## Materials and methods

### Serum samples

Serum samples were collected from subjects with diagnosis of SARS-CoV-2 infection confirmed by real-time RT-PCR testing of nasopharyngeal swabs. The presence and titer of SARS-CoV-2-neutralizing antibodies in serum was determined by an in-house virus microneutralization assay, which is considered the reference test in coronavirus serology. Disease severity was classified as (1) mild, non-hospitalized, (2) moderate, hospitalized, and (3) severe, admitted to the intensive care unit. Study subjects included 92 patients with mild or moderate COVID-19 and 15 patients with severe COVID-19. Leftover serum samples collected from 50 healthy subjects before December 2018 were used as control.

Specimens were stored at −20°C until use. The study was approved by the ethic committee of IRCCS Sacro Cuore Don Calabria Hospital and study subjects provided written informed consent.

### Proteome-wide microarray

A PEPperCHIP® Peptide Microarrays slides with 1 subarray, was used where each peptide sequence is composed by 15 aminoacids with an overlap of 13 residues. Serum samples were incubated dynamically over night at 4°C, diluted 1:50 in incubation buffer and subsequent protocol steps performed according to manufacturer’s instructions. Secondary antibody (Anti-Human IgG-Cy3, Jackson ImmunoReserarch) was used for fluorescence detection and microarrays scanned by TECAN power scanner at 10% laser intensity and 50% gain.

### Peptides synthesis

All peptides were synthesized by stepwise microwave-assisted Fmoc-SPPS on a Biotage ALSTRA Initiator+ peptide synthesizer according to well-established protocols. Briefly, peptides were assembled on a 2-CTC resin (0.5mmol/g loading) swelled in DCM/NMP 3:1. Chain elongation was performed by iterative cycles of amino acids coupling (using Oxyma/DIC as activators) and Fmoc-deprotection using a 20% piperidine solution in DMF. Coupling steps were performed for 20 minutes at 50°C. Fmoc-deprotection steps were performed by treatment with 20% piperidine solution at 40°C for 10 minutes. Following each coupling and deprotection step, peptidyl-resin was washed with DMF. Upon complete chain assembly, peptides were cleaved from the resin using a 2.5% TIS, 2.5% thioanisole, 2.5% water, 92.5% TFA mixture. Crude peptides were then purified by preparative RP-HPLC (Shimadzu). Finally, purified peptides were lyophilized and stored at −20°C.

### Peptide microarray

Silicon slides (SVM, Sunnyvail, CA) were coated by MCP6 (Lucidant Polymers) as follows: slides were treated with oxygen plasma for 15 min. MCP6 was dissolved in DI water to final concentration of 2% w/v and then diluted with ammonium sulphate solution 40% at ratio 1:1. Subsequently, silicon slides were immersed into the polymer solution for 30 min, then washed with DI water, dried under nitrogen steam and cured for 15 min under vacuum at 80°C.

Peptides were first dissolved in DMSO to 1 mM stock solution and then diluted to the final spotting concentration of 50μM in printing buffer composed by 25 mM Na/Acetate pH 4.8, 15 mM trehalose, CuSO_4_ 100 μM, ascorbic acid 6.25 μM and THPTA 400 μM. Slides were spotted using a non-contact Spotter S12 (Scienion Co., Berlin, Germany), in each subarray we used as positive control a peptide sequence derived from poliomyelitis virus. Printed slides were placed in a humid chamber overnight at room temperature. Then they were blocked in 2 mM EDTA water solution for 1 h, washed with water, and dried under a stream of nitrogen.

### Bioassay

All samples were diluted 1:50 in incubation buffer with 1% BSA, and incubated dynamically for 1 h at room temperature. Then, slides were washed 3 times, for 1 min. with washing buffer. The second incubation were performed with secondary antibody (Anti-Human IgG-Cy3, Jackson ImmunoReserarch; Anti-Human IgM-Cy5, Invitrogen) diluted in ratio 1:1000 in incubation buffer with 1% BSA. Finally, slides were washed and dried and the analysis were performed by TECAN power scanner 10% laser intensity and 50% gain.

The recombinant N full antigen for immunoreactivity testing was a kind gift from LifeLineLab, (Pomezia, Italy). The protein was immobilized on copoly(DMA-NAS-MAPS) coated slides and tested according to previously developed protocols ^15,16^

## Acknowledgements

Work partially funded by Regione Lombardia, project READY (Regional Network for developing diagnostic methods in rapid response to emerging epidemics and bio-emergencies) ID 229472, and by the European Union’s Horizon 2020 research and innovation programme, under grant agreement no. 874735 (project VEO).

