## Supplementary information for "SARS-CoV-2 epitope mapping on microarrays highlights strong immune-response to N protein region"

1: Consiglio Nazionale delle Ricerche, Istituto di Scienze e Tecnologie Chimiche “Giulio Natta”” (SCITEC), Via Mario Bianco 9, 20131, Milano, Italy

2: Department of Molecular Medicine, University of Padova, Via A. Gabelli 63, 35121 Padova, Italy

3: Department of Infectious-Tropical Diseases and Microbiology, IRCCS Sacro Cuore Don Calabria Hospital, Via Don A. Sempreboni, 5 – 37024 Negrar di Valpolicella (Verona), Italy

§: these authors equally contributed

#: these authors equally contributed

*: corresponding authors

**Supplementary information**

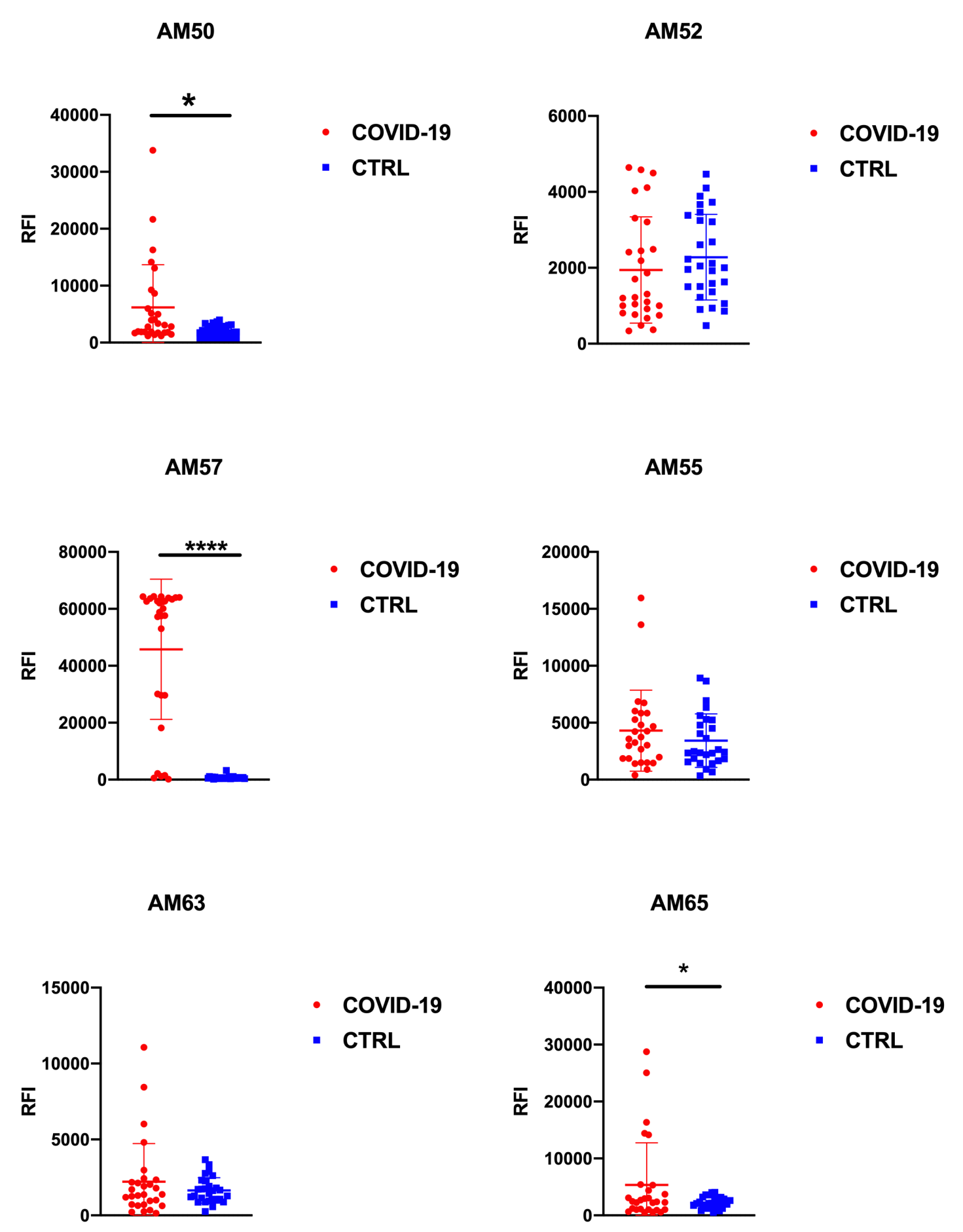

Figure 1: unpaired t-Test results for the peptide specific IgG. Peptide arrays displaying six peptide probes were probed with sera (N = 28) from COVID-19 patients and control patients (N = 28). Significative: p<0.05; * = p<0.05; ** = p<0.01; *** = p<0.001; **** = p<0.0001

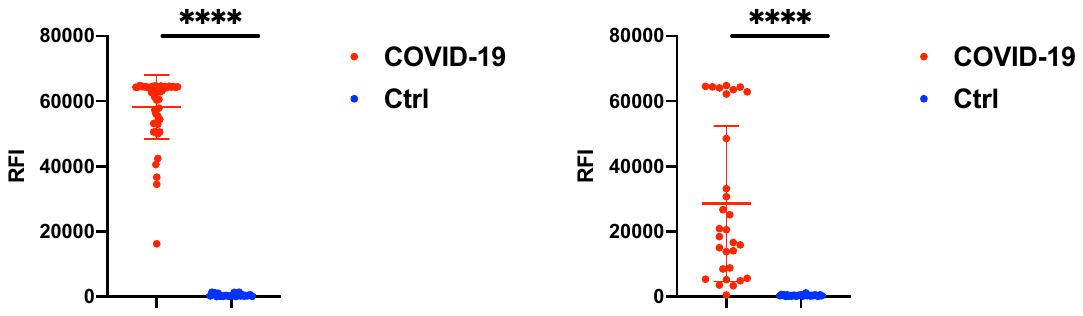

Figure 2. Left panel: IgG immunoreactivity on full N antigen. Right panel: IgM immunoreactivity on the full N antigen.
